# T Cell Predominant Response to AAV-Spike Protects hACE2 Mice from SARS-CoV-2 Pneumonia

**DOI:** 10.1101/2021.08.16.456441

**Authors:** Christopher D. Greer, Coral M. Kasden, Leon Morales, Kendall A. Lundgreen, Philip D. Hicks, Lilia J. Carpenter, Cristhian Salas-Quinchucua, Elisia D. Tichy, Albert M. Maguire, Jennifer G. Hoffman, Robin J. Bailey, Shangzhen Zhou, Angela Luo, Steven J. Chomistek, Benjamin W. Kozyak, Charles R. Bridges, Gilad S. Gordon, Geoffrey C. Tabin, Paul F. Bates, Jorge E. Osorio, Jean Bennett, Hansell H. Stedman

## Abstract

Prevention of COVID-19 is widely believed to depend on neutralization of SARS-CoV-2 by vaccine-induced humoral immunity^1,2^, raising concern that emerging escape variants may perpetuate the pandemic^3–6^. Here we show that a single intramuscular injection of Adeno-Associated Virus-6 (AAV6) or AAV9 encoding a modified, N-terminal domain deleted (ΔNTD) spike protein induces robust cellular immunity and provides long-term protection in k18-hACE2 transgenic mice from lethal SARS-CoV-2 challenge, associated weight loss and pneumonia independent of vaccine-induced neutralizing humoral immunity. In both mice and macaques, vaccine-induced cellular immunity results in the clearance of transduced muscle fibers coincident with macrophage and CD8+ cytotoxic T cell infiltration at the site of immunization. Additionally, mice demonstrate a strong Type-1 polarized cellular immunophenotype and equivalent *ex vivo* T cell reactivity to peptides of wt and alpha (B.1.1.7) variant spike. These studies demonstrate not only that AAV6 and AAV9 can function as effective vaccine platforms, but also that vaccines can provide long-term efficacy primarily through the induction of cellular immunity. The findings may provide an alternative approach to containment of the evolving COVID-19 pandemic and have broader implications for the development of variant-agnostic universal vaccines against a wider range of pathogens.

## Introduction

Spike protein is a class I homotrimeric fusion glycoprotein that initiates viral entry of host cells via high-affinity engagement of the membrane-bound receptor angiotensin-converting enzyme 2 (ACE2)^7^. Structure-guided mapping of immunodominant sites elicited by SARS-CoV-2 infection in COVID-19 patients has revealed that over 90% of neutralizing humoral responses are directed against antigenic sites in the ACE2 Receptor Binding Domain (RBD) of spike^8^. Because of spike’s essential function, it is widely assumed that vaccine efficacy will be tightly linked to titers of measurable neutralizing antibodies^1,2^. However, it has previously been suggested that T cell immunity alone might be sufficient to prevent COVID-19 based on the demonstration that cytotoxic CD8 T cells can protect mice from lethal challenge with mouse-adapted SARS-CoV following infusion of peptide-loaded dendritic cells^9^.

Adeno-associated viruses (AAVs) are a class of replication-deficient, non-pathogenic DNA viruses. AAVs have therefore become a preferred vector for clinical *in vivo* gene delivery and accordingly were the first vectors approved by the EMA and FDA for efficient local and systemic gene delivery in human genetic disease^10,11^. AAVs have single-stranded genomes of ^∼^4.8 kb, constraining the use of this vector system to synthetic genomes of <5kb. Based on our recent demonstration of robust T cell immunity against an AAV-delivered miniaturized dystrophin transgene in a canine model of muscular dystrophy^12^, we asked whether such an immune response to AAV-spike could be sufficient to protect against SARS-CoV-2.

### Design and Characterization of AAV Vaccine

The original Wuhan strain (HB-01) spike protein is encoded by a 3,822-nucleotide open reading frame in the RNA genome^13^. To accommodate the powerful and constitutive 832 bp CMV promoter used in our previous studies^12^, we designed two AAV-deliverable transgenes encoding mutant spike derivatives that selectively lack the autonomously folding N-terminal domain (ΔNTD) (Fig 1a). Both transgenes contain 8 total mutational sites and are therefore designated “M8” and “M8B,” with 7 of the 8 mutations being identical between both transgenes. In addition to two N-terminal deletions (Δ1 and Δ2) that together constitute ΔNTD, both transgenes share Q271G to afford structural flexibility immediately following Δ2, mutated furin and S2’ proteolytic cleavage sites, proline mutations to stabilize prefusion structure, and deletion of the C-terminal ER-retention signal (Δ3) (Fig. 1a)^14–19^. The alternative 8^th^ mutation was designed to selectively disrupt a conserved epitope encompassing the AA 614 position previously implicated in both antibody-dependent enhancement (ADE) in SARS-CoV and enhanced transmissibility of SARS-CoV-2^20–23^. The packaged genomes consist of codon-optimized M8 and M8B open reading frames in an AAV ITR-flanked transcriptional cassette identical to that previously described^12^ (Fig. 1b). We detected robust expression of M8 and M8B by western blot following transfection of HEK293 cells with plasmids containing the entire AAV-M8 and AAV-M8B genomes (Fig. 1c). M8B expression was additionally observed by immunofluorescent microscopy in transfected myogenic c2c12 cells (Extended Data Fig. 1).

**Figure 1:**
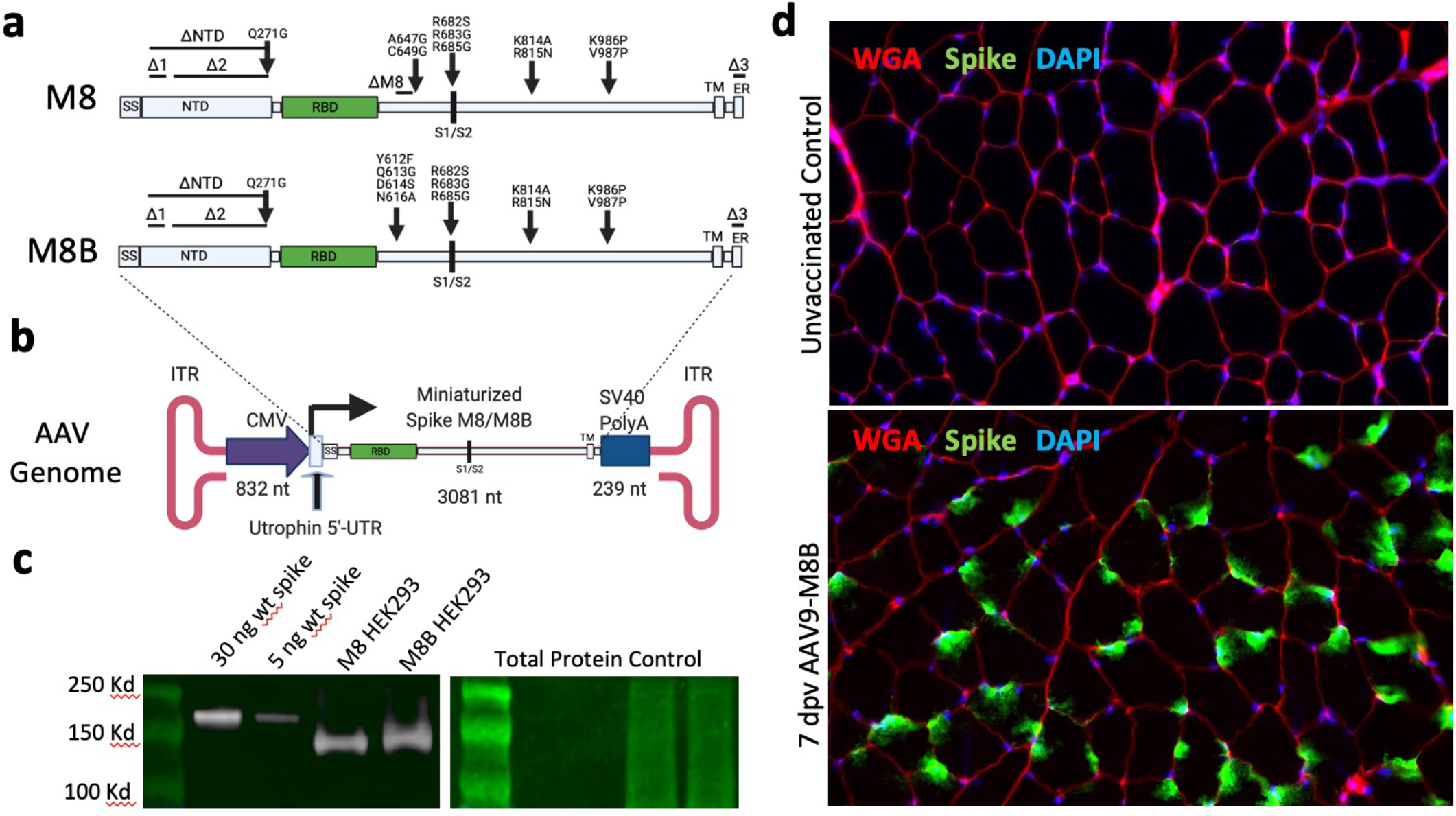
AAV vaccine design and characterization. **a,** Linear schematic of mutations and deletions (D’s) present in vaccine candidates M8 and M8B mapped onto spike protein from SARS-CoV-2 strain Wuhan/IVDC-HB-01/2019. SS: Signal sequence. NTD: N-Terminal Domain. RBD: Receptor Binding Domain. TM: Transmembrane Domain. ER: ER Retention Signal. **b,** Schematic depiction of recombinant AAV genome encoding a CMV promoter driven, codon-optimized, truncated spike transgene. **c,** Detection of M8 and M8B by western blot in HEK293 cells transfected with ITR-containing pAAVITRCMV-M8(b) plasmid. Recombinant spike protein was used as a positive control. **d,** Immunohistochemistry of C57BL/10 gastrocnemius muscle 7 days post IM vaccination of 6.4E10 vg AAV9-M8B (n=2) as well as an unvaccinated negative control. Wheat germ agglutinin (WGA) demarcates the muscles fiber membranes.

### Vaccine-Induced Immunity in C57BL/10 Mice

M8B was packaged into AAV9 (AAV9-M8B) and a group of eight C57BL/10 mice were administered 6.4E10 total viral genomes (vg) by an intramuscular (IM) route. Groups of two mice were euthanized at weekly intervals and immunohistochemistry was performed to assess the timecourse of M8B antigen expression as well as the overall muscle histology at the site of vaccination. At 7 days post vaccine administration (dpa) there was robust M8B expression within skeletal muscle fibers, while the muscle histology was otherwise normal (Fig. 1d). This finding remained unchanged until 21 dpa when regions of dense mononuclear infiltrates surrounded M8B-expressing muscle fibers. Immunohistological staining revealed F4/80+ macrophages (Fig. 2a) and CD8+ cytotoxic T cells (Fig. 2b) were present within the mononuclear infiltrates (Fig. 2a, b inserts), indicating an ongoing cellular immune response targeting the transduced muscle fibers. Actively regenerating, centrally nucleated muscle fibers resembling those in murine muscular dystrophy are seen in close proximity to these infiltrating immune cells, further suggesting immune-mediated elimination of transduced muscle fibers^12^ (Fig. 2b arrows). To control for immune response directed against the AAV9 vector capsid, we injected C57BL/10 mice with 6.4E10 vg AAV9-μUtrophin, a non-immunogenic vector in development for treatment of Duchenne muscular dystrophy^12^. At 21 dpa we observed the absence of macrophage or cytotoxic T cell infiltration, despite intense focal expression of the non-immunogenic μUtrophin transgene product (Extended Data Fig. 2).

**Figure 2:**
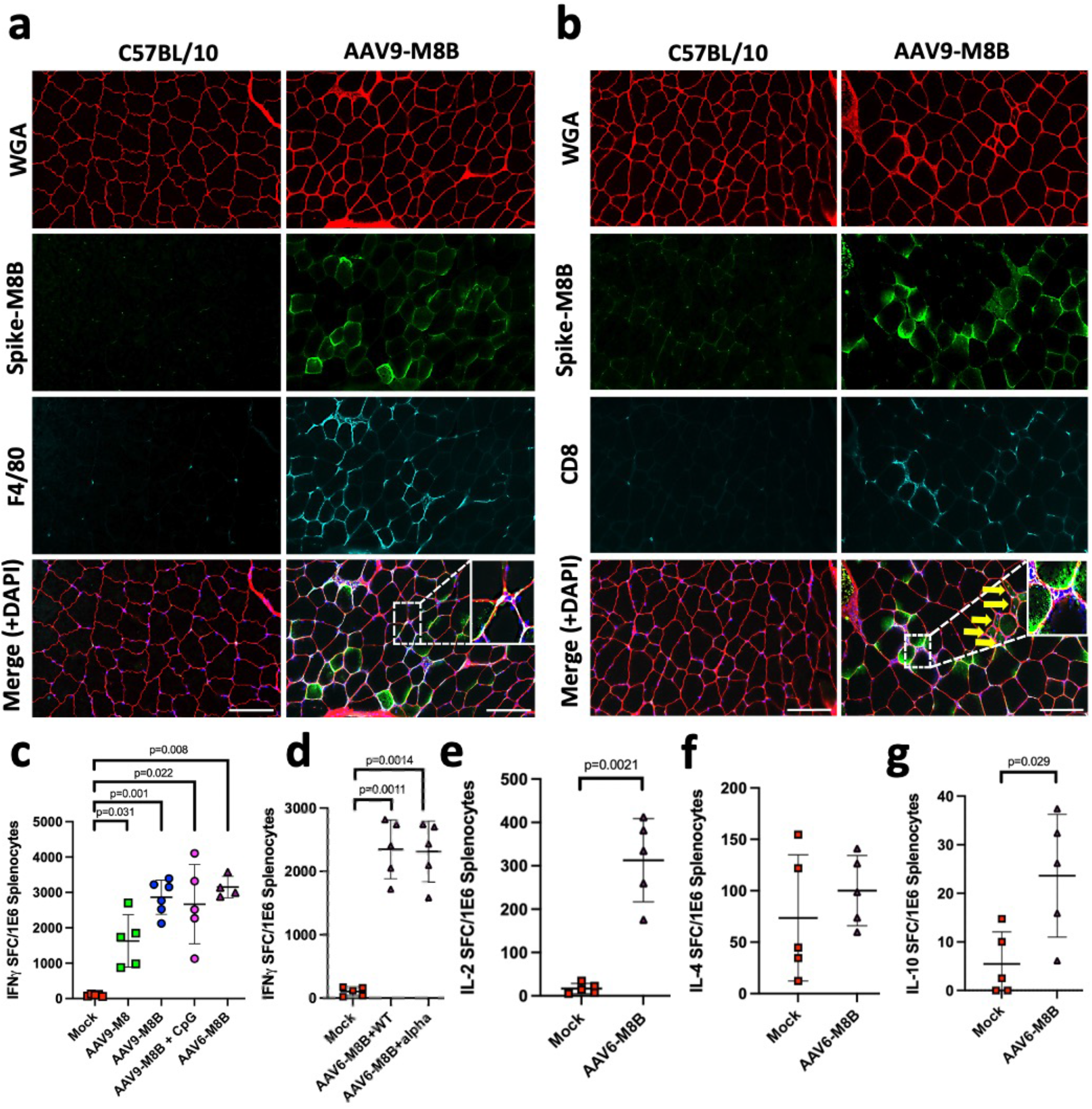
AAV Vaccine-Induced Cellular Immunogenicity in Mice. **a, b,** IHC shows F4/80+ macrophage (a) and CD8+ cytotoxic T-cell (b) infiltration surrounding M8B expressing muscle fibers at 3 weeks post IM injection of 6.4E10 vg AAV9-M8B (n=2). Scale bar = 100um. Yellow arrows in the merged images indicate centrally nucleated, actively regenerating muscle fibers (**b**). **c,** C57BL/10 mice received 6.4E10 vg IM injection of either AAV9-M8, AAV9-M8B, AAV6-M8B, or AAV buffer (PBS + 0.001% PF68) as a mock vaccinated control. At one month post vaccination, the indicated number of IFNγ secreting cells per 1E6 splenocytes were detected by ELISpot in response to *ex vivo* stimulation with peptide pools spanning both the S1 and S2 fragment of WT HB-01 spike. **d-g,** Mixed gender 2–3-month-old mice received either 1E12 vg AAV6-M8B (n=5 k18-hACE2 mice) or AAV buffer (n=2 k18-hACE2, n=3 C57BL/10). At 65 days post vaccination, the indicated number of IFNγ (**d**), IL-2 (**e**), IL-4 (**f**), and IL-10 (**g**) secreting cells per 1E6 splenocytes were detected by ELISpot in response to *ex vivo* stimulation with individual peptide pools spanning the S1 and S2 fragment of WT HB-01 spike. Alpha variant spike peptide pools were used in **d** where indicated. **(c-g)** Symbols represent individual animals. Lines show mean +/− 1 S.D. Results were compared either by Brown–Forsythe and Welch’s analysis of variance (ANOVA) with Dunnett’s T3 multiple comparisons test (**c, d**) or Welch’s t-test (**e-g**).

To characterize the T cell responses, C57BL/10 mice received a single IM administration of either 6.4E10 vg AAV9-M8, AAV9-M8B, or AAV6-M8B and splenocytes were harvested one month post administration (mpa). Following ex vivo stimulation with peptide pools spanning the S1 and S2 fragment of spike, enzyme-linked immune absorbent spot (ELISpot) assays demonstrated many IFNγ-secreting cells in all vaccine groups (Fig. 2c), evidence of a strong Type 1 immune response. These data indicate that a single IM dose of AAV-encoding either M8 or M8B is capable of eliciting a robust T cell mediated immune response against the AAV-encoded transgene product *in vivo*.

### Immunity in and Protection of k18-hACE2 Mice

Angiotensin-converting enzyme 2 (ACE2) is the receptor for SARS-CoV-2 in humans^13^. Wild type mice are not susceptible to SARS-CoV-2 infection because the spike RBD has low affinity to the mouse ACE2 ortholog. k18-hACE2 transgenic mice express human ACE2 (hACE2) driven by the cytokeratin 18 (k18) promoter in airway and other epithelial cells^24^. It was recently demonstrated that inoculation of this strain with 10^4 plaque-forming units (PFU) generates 50% lethal respiratory disease resembling severe COVID-19 [^24,25^.

To characterize the vaccine-induced memory response in k18-hACE2 mice, animals received a single IM injection of either 1E12 vg AAV6-M8B or AAV buffer and splenocytes were harvested at 65 dpa. ELISpot revealed significant numbers of splenocytes secreted T-helper-1 (Th1)-type cytokines IFNγ and IL-2 upon *ex vivo* stimulation with peptide pools spanning the S1 and S2 fragment of spike (Fig. 2d, e). Interestingly, at 65 dpa there is no significant increase in number of splenocytes secreting the canonical T-helper-2 (Th2)-type cytokine IL-4 upon identical *ex vivo* stimulation (Fig. 2f); however, a significant, albeit very small total number of cells secrete IL-10 (Fig. 2g). Finally, to assess whether T cell responses are maintained in response to exposure to spike proteins from variant SARS-CoV-2 strains, we ran a second IFNγ ELISpot and stimulated *ex vivo* with peptide pools spanning the S1 and S2 fragments of spike protein of the alpha variant and there was no change in the number of IFNγ secreting splenocytes (Fig. 2d). Together, these experiments indicate that the cellular memory response against the spike is strongly Type 1-polarized, and this response is not diminished by mutations present in the alpha variant spike.

To assess vaccine efficacy against lethal SARS-CoV-2 challenge and associated viral pneumonia, 6–8-week-old k18-hACE2 mice were administered a single dose IM injection of either 1E12 vg AAV9-M8, AAV9-M8B, AAV6-M8B, or AAV buffer as a mock vaccinated control group before intranasal inoculation with a lethal dose of 2.5×10^4 PFU of SARS-CoV-2 at 25 dpa (Extended Data Fig. 3a). In order to compare tissues and viral RNA levels between groups, we decided to terminate the experiment at 7 days post infection (dpi) with SARS-CoV-2, by which time the majority of k18-hACE2 mice will have died or met objective criteria for euthanasia by the masked/blinded veterinarians overseeing the challenge experiments. As expected, by 6 dpi 100% of mock-vaccinated animals met terminal euthanasia criteria as determined by the presence of all of the following clinical findings: absence of movement, persistent eye closure with palpebral crusting, nasal secretions, kyphosis, and the presence of a rough coat resembling mice with the “rc” mutation^26^. No vaccinated animal demonstrated any of these clinical symptoms at any point in the 7-day experiment (Extended Data Fig. 3b).

While the experimental design (termination at 7 dpi) did not allow for a longer assessment of differential survival, it did permit assessment of other objective surrogates of vaccine-induced protection. Mock-vaccinated mice began progressively losing weight starting at 4 dpi whereas animals in the vaccinated groups maintained stable weight over the 7 days post challenge (Fig. 3a, Extended Data Fig. 3c). Animals were assessed for cardiopulmonary performance at 5 dpi by measuring the distance they were able to run on a treadmill for 8 minutes according to a predetermined ramping protocol. Whereas 80% of the mock vaccinated mice performed worse on the treadmill at 5 dpi than they did a week prior to viral challenge, all mice in each vaccinated group either improved or had no change in performance (Fig. 3b, Extended Data Fig. 3d). Respiratory performance of these animals was assessed by whole body plethysmography at 5 and 6 dpi and there was a significant drop off in both tidal volume and respiratory rate in the mock vaccinated group at each of these time points relative to the pre-challenge assessment of all k18-hACE2 mice (Fig. 3c, d). Consistent with the observed differences in exercise tolerance and respiration, lungs of AAV-injected mice appeared grossly normal, while lungs of PBS-injected mice revealed signs of severe interstitial pneumonia characterized by collapse of the alveolar spaces (Fig. 3e). We quantified viral RNA present in the lungs and observed a 2.85-log reduction for AAV9-M8B, a 4.44-log reduction for AAV9-M8, and a 4.3-log reduction for AA6-M8B vaccinated relative to the mock-vaccinated control group (Fig. 3f), consistent with the significant decrease in spike protein in lung tissue sections by immunohistochemistry (Extended Data Fig. 3e).

**Figure 3:**
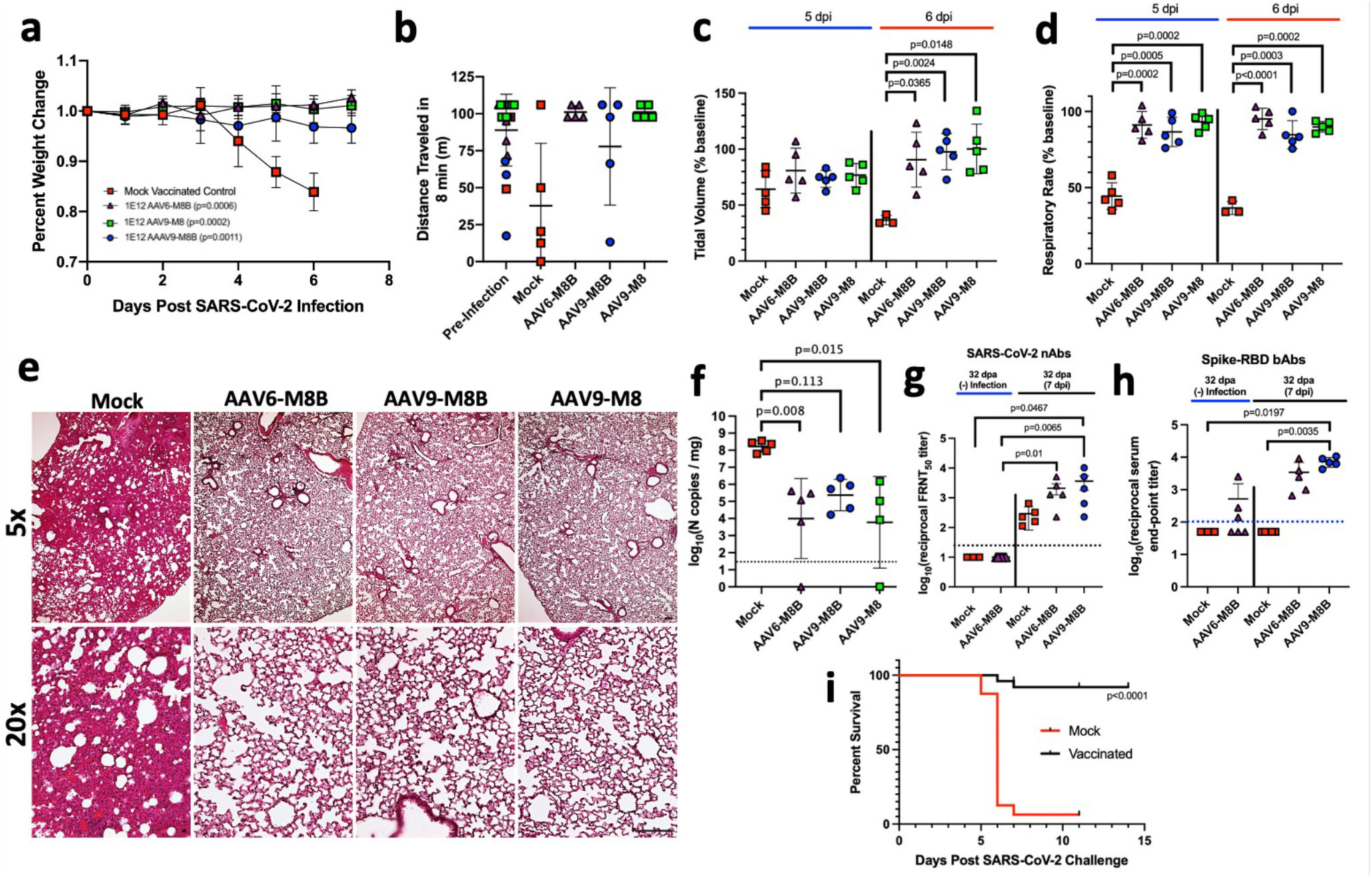
AAV-M8/M8B protects transgenic k18-hACE2 mice from lethal SARS-CoV-2 challenge and associated severe disease. **a-i,** k18-hACE2 mice received an IM injection of either 1E12 vg AAV6-M8B (n=5), 1E12 vg AAV9-M8B (n=5), 1E12 AAV9-M8 (n=5), or AAV formulation buffer (n=5) as a mock vaccination control. Genders within groups were randomized with either 2 males/3 females, or the inverse. **a,** Weight trace post challenge. Groups were compared by two-way ANOVA with Dunnett’s multiple comparisons test. p-values shown in figure are at 6 dpi compared to mock. At 5 dpi, p<0.01 for AAV9-M8b and p<0.001 for AAV6-M8B and AAV9-M8. **b,** Results of a treadmill performance test at 5 dpi, as measured by distance traveled in 8 min. **c, d,** Results of the respiratory assessment at both 5 and 6 dpi, as measured by tidal volume (**c**) and respiratory rate (**d**). Days 5 and 6 were independently compared by Brown–Forsythe and Welch’s analysis of variance (ANOVA) with Dunnett’s T3 multiple comparisons test. **e**, Representative lung histopathology. Scale bar = 100um. **f,** Viral RNA quantified from whole lung homogenates. **g,** Anti-Spike RBD IgG titers determined by ELISA following viral challenge compared to a group of 1E12 vg AAV6-M8B vaccinated, but unchallenged mice (n=6) at the equivalent timepoint post-vaccination. **h,** SARS-CoV-2 neutralizing antibody titers determined by focus reduction neutralization test (FRNT_50_). This was similarly compared to the unchallenged AAV6-M8B vaccinated group as in (**g**). **h,** Kaplan Meier curve depicting aggregate survival for all unvaccinated and 1E12 vg AAV vaccinated k18-hACE2 mice post infection with 2.5E4 PFU SARS-CoV-2 (AAV6-M8B n=15, AAV9-M8B n=5, AAV9-M8 n=5). Data generated from 3 independent experiments. Survival was compared by the log-ranked Mantel-Cox test. (**b-d**) Each shape indicates an individual mouse, and the lines show the mean +/− 1 S.D. (**g**) Symbols represent the geometric mean for all replicates of an individual animal. (**f-h**) the dotted line indicates the limit of detection, and the lines show the mean +/− 1 S.D. Groups were compared by Kruskal–Wallis analysis of variance (ANOVA) with Dunn’s multiple comparisons test

Interestingly, our analysis of serum obtained at necropsy revealed that this robust protection occurred despite the absence of vaccine-induced neutralizing humoral immunity (Fig. 3g). This finding was confirmed in mice by giving either AAV9-M8 or AAV9-M8B followed by a booster 5 weeks later. Three weeks post booster there was still an absence of detectable neutralizing antibody (Extended Data Fig. 4). However, to support the claim AAV-M8B-primed CD4+ T cells enhance the kinetics of neutralizing antibody production post-SARS-CoV-2 infection, we show that neutralizing antibodies in the challenged AAV6-M8B and AAV9-M8B groups, but not the challenged mock vaccinated group, are significantly elevated relative to the unchallenged AAV6-M8B vaccinated group at the completion of the experiment (Fig 3g). To further support the claim of accelerated post-vaccination, post-infection humoral immunity kinetics, we assayed for IgG titers against recombinant spike RBD. Some of the unchallenged AAV6-M8B group did have detectable, albeit low levels, of anti-RBD IgG whereas all the challenged mock vaccinated animals were below the limit of detection (Fig. 3h). However, the challenged AAV6-M8B and AAV9-M8B vaccinated mice were elevated 6.6-fold and 14.3-fold, respectively, relative to the unchallenged AAV6-M8B group and the challenged AAV9-M8B groups RBD IgG titer was significantly greater than that of the challenged mock vaccinated group (Fig. 3h). Together, these serological studies reveal that AAV-M8/M8B vaccination alone produces a very mild humoral immune response with no detectable neutralizing antibody component; however, upon infection vaccinated mice respond with an accelerated humoral response.

We next assessed long-term therapeutic efficacy as well as vaccine stability in real-world storage conditions, specifically following long-term refrigeration. A cryofrozen vial of AAV6-M8B was thawed and used to vaccinate k18-hACE2 mice (Cohort 1). The AAV6-M8B vial was then stored at 4°C for 4 months before being used to vaccinate a second group of k18-hACE2 mice (Cohort 2) and then both cohorts were simultaneously challenged with 2.5×10^4 PFU SARS-CoV-2 IN ^∼^18 days later (Extended data Fig. 5a). We observed long-term protection from lethality and weight loss 4.5 months out from single dose vaccination in Cohort 1 (Extended Data Fig. 5b-d). Furthermore, long-term refrigeration did not impair its ability to protect against lethal challenge in Cohort 2 even at only 18 dpa (Extended Data Fig. 5e); however, protection against weight loss was not statistically significant at this early timepoint post vaccination (Extended data Fig. 5f, g).

In aggregate, these challenge experiments demonstrate that vaccination with AAV-M8/M8B provides a highly significant survival advantage to k18-hACE2 mice upon subsequent lethal challenge with SARS-CoV-2 (Fig. 3).

### Vaccine-Induced Immunity in Macaques

To preliminarily assess capsid efficacy and dose-response in non-human primates, four cynomolgus macaques were vaccinated IM with either AAV6-M8B or AAV9-M8B at a dose of either 1E11 vg or 1E12 vg. At 29 dpa there is significant mononuclear cell infiltration and the presence of cytotoxic CD8 T cells and CD11b+ macrophages surrounding M8B expressing muscle fibers (Fig. 4a, b). IFNγ ELISpots performed on peripheral blood mononuclear cells (PBMCs) isolated at 29 dpa demonstrated the presence of spike antigen-specific T cells in both macaques that received the higher dose of 1E12 vg, but neither animal that received the lower dose of 1E11 vg (Fig. 4c). Finally, as observed in mice, there was a small increase in the anti-Spike-ECD IgG titers of both macaques that received the 1E12 vg dose, but neither animal that received the dose of 1E11 vg and no vaccine-induced neutralizing antibodies were detectable at 29 dpa at any dose (Fig. 4d, e). These data strongly support our data generated from mice and suggest a similar immune response to AAV-M8B in primates.

**Figure 4:**
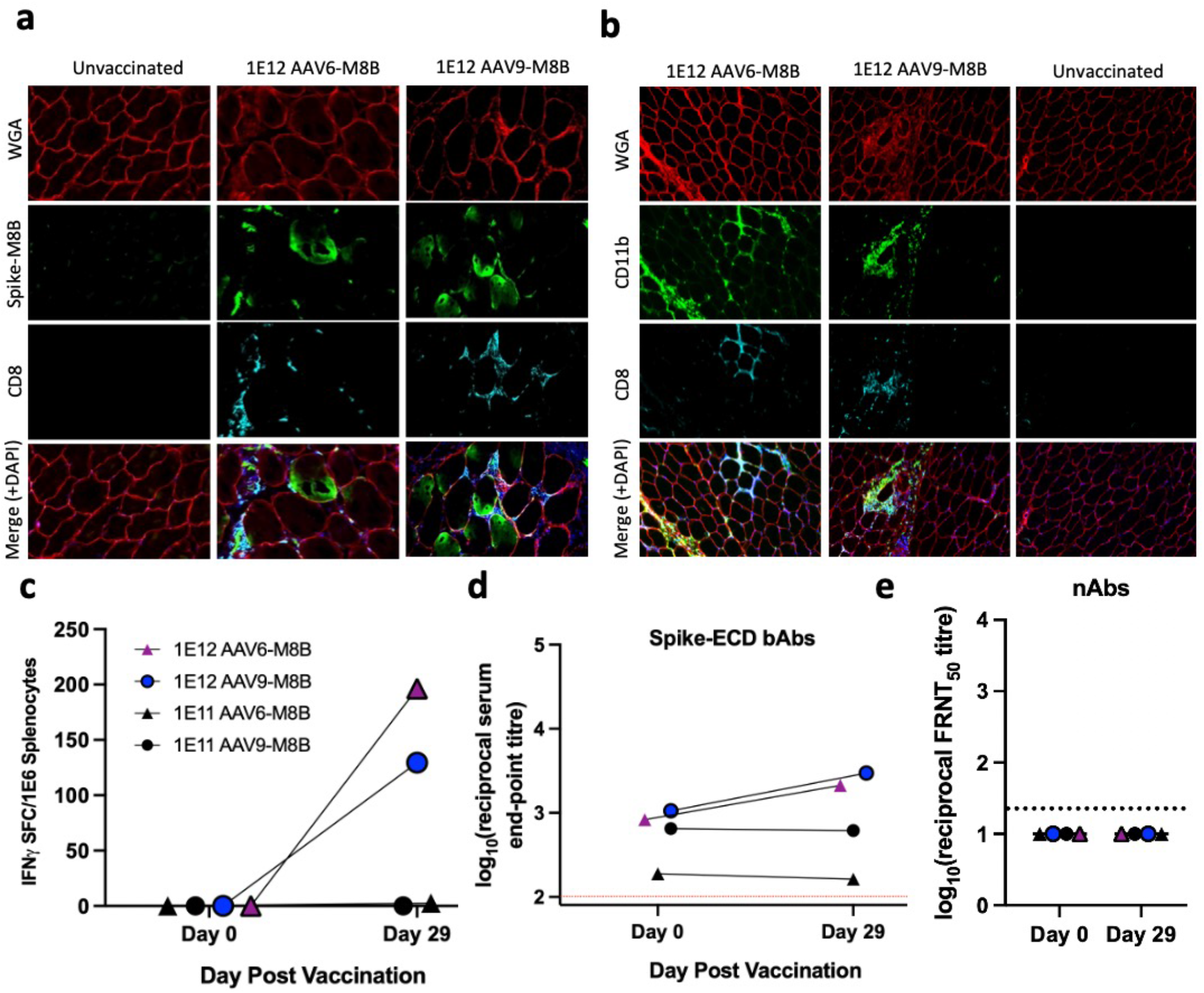
AAV Vaccine Immunogenicity in Macaques. **a,** IHC shows CD8+ cytotoxic T cell infiltration surrounding M8B expressing muscle fibers 29 days after a single IM vaccination with either 1E12 vg of AAV6-M8B or AAV9-M8B. **b,** IHC shows CD11b+ macrophages in the areas nearby CD8+ cytotoxic T cells surrounding muscle fibers following IM vaccination. **c,** Number of IFNγ secreting cells per 1E6 PBMCs as detected by ELISpot. **d,** Anti-Spike ECD IgG titers determined by ELISA. **e,** SARS-CoV-2 neutralizing antibody titers determined by focus reduction neutralization test (FRNT_50_).

## Discussion

Here we have shown that AAV vectors encoding N-terminally deleted spike proteins are capable of inducing robust, long-term protection of k18-hACE2 mice from lethal SARS-CoV-2 challenge as well as associated viral pneumonia. This protection in mice was mediated by a Type-1 polarized cellular immune response in the absence of a detectable neutralizing humoral immune response. We have also demonstrated that AAV vaccines remain effective even after months of standard refrigeration and are effective and durable following single dose administration. These features make the vaccine platform potentially advantageous for distribution to rural, sparsely populated, or resource-limited regions where long-term vaccine thermostability may be critical.

Interestingly, the observed protection against viral challenge occurred in the absence of vaccine-induced neutralizing antibodies, thereby formally demonstrating that neutralizing humoral immunity is not a prerequisite for protection against SARS-CoV-2 infection. In addition to providing immediate protection against viral replication in host cells, strong and durable T cell memory in recipients of an AAV-M8/M8B vaccine would be expected to accelerate affinity maturation of neutralizing antibodies specific for a mutant RBD in any SARS-CoV-2 variant to which the host is initially exposed. Our findings with regard to similar T cell reactivity to peptides from the alpha variant illustrate an example of such preservation of effector and helper T cell responses. A theoretical advantage of this variant-agnostic approach for a vaccinated population is its potential to circumvent the downside of antigenic imprinting to RBD conformational epitopes specific to the original Wuhan SARS-CoV-2 Spike. Among individuals in an outbred human population, the identity of immunodominant T cell epitopes will vary widely across the Spike sequence on the basis of HLA polymorphism, in contrast to the focal concentration of conformational B cell epitopes in the RBD to which most individuals must target neutralizing antibodies. Thus, the strength and durability of vaccine-induced T cell immunity has important implications for the future evolution of the COVID-19 pandemic as herd immunity to the original variant escalates and selective pressure builds for SARS-CoV-2 variants capable of escaping neutralizing humoral immunity through mutations in the RBD.

## Acknowledgements

This research was funded by 4MVac, LLC, the University of Pennsylvania Cardiovascular Institute, and the National Respiratory Distress Syndrome Foundation in memory of Timothy S. Cannon. We are grateful for veterinary support in the rodent and non-human primate studies, with particular thanks for the latter to Dr. Alicia Donnelly and team. We are grateful from support of the Center for Advanced Retinal and Ocular Therapeutics and its Research Vector Core for generation of the AAV used in this study. We also thank the UPenn Human Immunology Core, the UPenn Gene Therapy Program Immunology Core, and the CDB microscopy core for their technical services and in particularly Jessica Chichester, Andrea Stout, and Jasmine Zhao for their technical assistance.

## Author contributions

H.H.S. conceived of the project. C.D.G. and H.H.S. designed the experiments. C.D.G., C.K., L.M., K.L, P.D.H, L.J.C., C.S.Q., A.M., E.D.T., J.H., R.B., S.C., and J.B. performed experiments and interpreted data. S.Z. and A.L. generated required reagents. B.W.K., C.R.B., G.T., G.G., P.B., J.B., and J.O. contributed analysis and interpretation. C.D.G. wrote and H.H.S. edited the manuscript. C.D.G. and H.H.S. oversaw all aspects of the research.

## Methods

### M8/M8B transgene synthesis

*De novo* gene synthesis of M8 and M8B cDNA was performed by GeneArt using the codon optimization scheme from (Sino Biological, VG40589-UT). M8/M8B were cloned into the AAV transfer plasmid pZac2.1 along with (or flanked by) a CMV promoter and SV40 PolyA sequence. Cloned plasmids were transformed into Stbl2 competent cells and selected by ampicillin resistance.

### AAV6/9 vector generation

AAV6 and AAV9 were individually generated and purified by the University of Pennsylvania preclinical vector core using the triple transfection method in HEK293 cells as previously described^12^. Vector preparations were assayed for quality, purity, and endotoxin levels. Vector aliquots were stored at −80°C and thawed immediately prior to IM injection, except where explicitly stated for experimental purposes.

### Transfection and Immunocytochemistry of C2C12 cells

3.0E4 C2C12 cells were plated into each well of an 8 well chamber slide (Millipore, PEZGS0816) and cultured for 24 hours in growth media (DMEM high glucose supplemented with 10% FBS, 1× anti-anti, 1× GlutaMAX, 1× NEAA). Cells were then transfected with 0.4 ug plasmid DNA using lipofectamine 3000 in growth media for 5 hours at 37°C before being switched to C2C12 differentiation medium (DMDM high glucose, 5% horse serum, 1× anti-anti, 1× NEAA, 1× glutamax) for 72 hours. Cells were then fixed with 10% NBF for 10 min at room temperature and then washed for 5 minutes with PBS before being permeabilized in 0.5% Triton X-100 solution 10 minutes at room temp. Cells were washed in PBS for 5 min before being blocked with 10% normal donkey serum (Abcam, ab7475, Lot GR3234297-32) diluted in PBS for 20 minutes at room temperature, followed by incubation with rabbit polyclonal a-SARS-CoV-2 Spike/RBD (Sino 40592-T62, Lot HD14AU0606 diluted 1:50 in PBS) for 1 hour at 37°C. Slides were then washed 3×10 minutes in PBS at RT and again blocked with 10% normal donkey serum (Abcam, ab7475, Lot GR3234297-32) for 20 minutes at room temperature before being incubated with PBS containing both WGA-555 (Thermo Fisher, W32464, diluted 1:40) and donkey a-rabbit-AF488 (Abcam, ab150061, diluted 1:500) for 30 minutes at 37°C. Following another 3×10 min washes in PBS, slides were mounted in VECTASHIELD mounting media containing DAPI (Vector Laboratories, H-1500). Images were taken using a Zeiss Observer 7 widefield microscope (Zeiss).

### Western Blot of transfected HEK293

M8/M8B-pZac2.1 plasmid transfected HEK293 cell pellets were lysed in RIPA lysis buffer (Santa Cruz Biotechnology, sc-24948) supplemented with protease inhibitor cocktail (Roche, 39802300). After homogenization, samples were spun at max speed for 15 minutes at 4°C and the supernatant was then transferred to a new 1.5 ml microcentrifuge tube. Protein concentration was determined for each sample using the Pierce BCA protein assay kit (Thermo Scientific, 23227). Samples were then diluted to an appropriate concentration with 6× Laemmli SDS sample buffer (Alfa Aesar, J61337) before heat denaturing for 5 minutes at 95°C. 30 ⍰g total protein was loaded per lane after transfer to a nitrocellulose membrane total protein was imaged for loading control using the REVERT Total Protein Stain Kit (LI-COR, 926-11010) and imaged by 700 nm fluorescence using the Odyssey Fc infrared imaging system (LI-COR). The membrane was then stained overnight at 4° in PBST + 5% BSA containing anti-SARS-CoV-2 Spike/RBD (Sino 40592-T62, Lot HD14AU0606 diluted 1:1500). The membrane was then washed 3×15 minutes in PBST and then incubated in secondary antibody solution (PBST + 5% BSA + Goat a-Rabbit-HRP 1:5000) (Abcam, ab205718) for one hour at room temperature. The membrane was then washed again 3×15 minutes in PBST before visualizing on the Odyssey Fc imaging system (LI-COR) using SuperSignal West Pico PLUS Chemiluminescent Substrate (Thermo Scientific, 34580).

### Mice

Murine experiments were executed in compliance with approval from the Animal Care and Use Committees of the University of Pennsylvania and University of Wisconsin at Madison. c57BL/10SnJ and B6.Cg-Tg(K18-ACE2)2Prlmn/J mice were purchased from the Jackson Laboratory.

### Vaccination of c57BL/10 and k18-hACE2 mice

Immediately prior to IM injection, the AAV was removed from the −80°C freezer and thawed at room temperature. The AAV vector was then diluted in formulation buffer (PBS + 0.001% Pluronic F-68) such that 50 ul of diluted vector contained the desired dose of viral genomes (either 6.4E10 or 1E12 vg), this controlled for volume injected across either dose administered. Mixed gender 2-5-month-old C57BL/10 mice or 6–9-week-old k18-hACE2 mice were randomly assigned to groups and anesthetized with isoflurane before receiving an intra gastrocnemius injection of either 50 ul AAV or 50 ul formulation buffer using a custom 100 uL Hamilton syringe with a 32-gauge needle (475-41182, Lot 81008). Additionally, k18-hACE2 mice were ear tagged as to keep the team performing the challenge experiments blinded to the animal’s treatment status.

### c57BL/10 muscle procurement, sectioning, and storage for immunohistochemistry

At pre-determined timepoints post-vaccination, mice were anesthetized with 4% isoflurane in an anesthesia chamber before being transferred to a nose cone where they were maintained on 4% isoflurane in 100% O_2_ at a flow rate of 1 L/minute. Toe pinch was performed to ensure mice were deeply anesthetized before mice were euthanized by cervical dislocation. Gastrocnemius muscles were harvested and embedded in O.C.T. (Tissue-Tek, 4583) within 7×7×5 mm base molds (Richard-Allan Scientific, 58949) by rapid submersion in liquid nitrogen-cooled isopentane. All OCT embedded tissue blocks were stored at −80 °C. 12-24 hours before sectioning, OCT blocks were moved to −20°C to gradually to warm to this temperature for optimal sectioning. Cryosections of 10-12 μm thickness were cut on a cryostat (Microm HM550, Thermo Scientific) at −22 °C and mounted on glass slides (Fisher Scientific, Superfrost Plus, 12-550-15). Slides were allowed to air dry for 10 minutes at room temperature before being stored at −20°C.

### c57BL/10 Immunohistochemistry

10-12 μm gastrocnemius cross-section containing slides were removed from −20°C and allowed to dry at room temperature for 10 minutes. Slides were then rehydrated for 10 minutes in PBS, permeabilized in 0.1% solution of Triton X-100 diluted in PBS for 10 minutes, and then washed again in PBS for 5 min. Sections were then blocked with 10% normal donkey serum (Abcam, ab7475, Lot GR3234297-32) diluted in PBS for 20 minutes at room temperature, followed by incubation with either rabbit polyclonal anti-Utrophin (custom, Sigma) or rabbit polyclonal anti-SARS-CoV-2 Spike/RBD (Sino, 40592-T62), with or without rat anti-CD8 (Thermo Fisher, MA5-17594) or rat anti-F4/80 (Thermo Fisher, MA1-91124), for 1 hour at 37°C. Slides were then washed 3×10 minutes in PBS at RT and again blocked with 10% normal donkey serum (Abcam, ab7475, Lot GR3234297-32) for 20 minutes at room temperature before being incubated with PBS containing WGA-555 (Thermo Fisher, W32464), donkey anti-rabbit-AF488 (Abcam, ab150061), and donkey anti-rat-AF647 (Abcam, ab150155) for 30 minutes at 37°C. Following another 3×10 min washes in PBS, slides were mounted in VECTASHIELD mounting media containing DAPI (Vector Laboratories, H-1500). Images were taken using a Zeiss Observer 7 widefield microscope (Zeiss).

### SARS-CoV-2 Viral stock

SARS-CoV-2 strain USA-WA1/2020 was provided by BEI (NR-52281), propagation was performed by passaging two times in Vero E6 cells in DMEM supplemented with 2% fetal bovine serum and 1× Antibiotic Antimycotic Solution (sigma A5955). Supernatant was collected upon observation of cytopathic effect and centrifuged for debris removal. Titration was performed in triplicate by plaque assay in 6 well plates with crystal violet dye using a 200μL inoculum per well from the generated stock. The produced stock with a final titer of 3×10^6^ PFU/ml was aliquoted and stored at −80°C.

### SARS-CoV-2 Challenge

All SARS-CoV-2 intranasal inoculations and subsequent physiological assessments and monitoring were done blinded to animal vaccination status. Infection of mice was conducted in a certified Animal Biosafety Level 3 (ABSL-3) laboratory at University of Wisconsin - Madison. The protocol for the challenge studies was approved by the University of Wisconsin Institutional Animal Care and Use Committee (Protocol Number V006324). Animals were anesthetized with 5% isoflurane, and 2.5×10^4^ PFU of SARS-CoV-2 (USA-WA1/2020) in a 50μL volume was administered via Intranasal inoculation using a micropipette tip. On the day of challenge and each day after animals were weighed and monitored for health issues, loss of weight, inappetence, activity reduction, and respiratory distress. Animals were euthanized by exposing them to an overdose of isoflurane in a closed chamber followed by cervical dislocation at the completion of the experiment or when presented with the following predetermined human endpoint criteria-- loss of greater than 20% body weight or the presence of all the following clinical signs: respiratory distress, inappetence, signs of dehydration, lack of mobility and critical body condition. Before euthanasia, blood serum was obtained by maxillary vein collection. After euthanasia, right lung for viral loads were collected, weighed, and homogenized in 1 mL of Trizol using a bead beater, after centrifugation, supernatant was stored at −80°C. Left lung was separated and inflated with 10% neutral buffered formalin using a 26G veterinary I.V. catheter and preserved in the formalin for histopathological changes evaluation.

### Treadmill acclimatization and performance assessment

Vaccinated and mock vaccinated k18-hACE2 mice were acclimated to treadmill twice prior to assessing their 8-minute treadmill performance 6 days before SARS-CoV-2 infection. These same mice were then assessed according to the same performance test at 5 days post SARS-CoV-2 infection. Treadmill performance tests followed a sex specific, 8-minute speed ramping protocol and the distance recorded was either that maximally allowed by the protocol in 8-minutes or that which they had completed upon their fifth 1.2 mA electrical shock.

### Whole body plethysmography

A custom whole body plethysmography system was designed for use in the ABSL3 facility at the University of Wisconsin to minimize both the need for mouse handling and the potential for mouse-to-mouse SARS-CoV-2 contamination. Pressure deflections in 500 cc single-use chambers were recorded into Acqknowledge 4.2 files using the Biopac TSD160A, DA100C, UIM100A, ad MP150 hardware modules in series. The DA100C module was configured to the following settings: GAIN 1000, 10HzLP ON, LP 5kHz, and HP 0.05Hz. Investigators blinded to group assignments recorded ear tag number of individual mice and then placed them into a chamber to enable a 60 second period of uninterrupted pressure recording. To minimize background noise, the raw signal collected was filtered via Acqknowledge 4.2 Software using the following settings: Low Pass sample rate at 8Hz, High Pass sample rate at 1.5Hz. Rate Signal parameter and identification of Peaks was determined using a threshold of 0.002 Volts. All files were then managed identically to enable extraction of average values and standard deviations for two parameters over 60 seconds: respiratory rate and amplitude, the latter proportional to relative tidal volume. The results for individual mice were normalized to baseline data obtained before inoculation with SARS-CoV-2. The sensitivity and specificity of the system was validated in mice recovering from inhalational general anesthesia using brief exposure to isoflurane at 4% of inspired gas.

### Lung Histology

Left lung tissues were fixed in 10% neutral buffered formalin then paraffin-embedded and sectioned at 7um. Sections were deparaffinized in two changes of xylene followed by rehydration through descending concentrations of ethanol (100%, 95%, 80%, water). Sections were stained in Gill’s hematoxylin (Thermo Scientific Cat# 72604) for 3 minutes, then rinsed under running tap water, differentiated in 0.1% acetic acid for 3 seconds, followed by a second rinse. Sections were then immersed in Scott’s bluing reagent (Ricca Cat# 6697-1) for 1 minute and transferred to water, then counterstained in eosin-Y solution (Thermo Fisher Cat# 71204) for 30 seconds. Finally, slides were dehydrated from water through ascending concentration of ethanol (70%, 95%, two changes 100%), cleared in two changes of xylene, and mounted with Cytoseal 60 (Thermo Scientific Cat# 8310-4) and coverslip. Sections were visualized by brightfield microscopy.

### SARS-CoV-2 viral load quantification

RNA was extracted from tissue homogenates using a Trizol/chloroform extraction method, and cDNA was synthesized using high-capacity cDNA to RNA-to-cDNA Kit (Thermo Scientific). SARS-CoV-2 RNA levels were measured by quantitative PCR using TaqMan Fast Universal PCR Master Mix (Thermo Fisher: Cat# 4352042) on the Applied Biosystems 7500 Fast Dx Real-Time PCR System. SARS-CoV-2-specific primers and probe set from the CDC 2019-Novel Coronavirus (2019-nCoV) Real-Time RT-PCR Diagnostic Panel were used to target the SARS-CoV-2 nucleocapsid (IDT: Cat.# 10006713). CoV_N1 F Primer: GACCCCAAAATCAGCGAAAT; CoV_N1 R primer: TCTGGTTACTGCCAGTTGAATCTG; CoV_N1 probe: FAM-ACCCCGCATTACGTTTGGTGGACC-BHQ1. A standard curve was generated with SARS-CoV-2 nCov_N_Positive control plasmid (IDT: Cat. #10006625) to determine viral copy numbers. Viral gene expression levels were normalized to lung tissue weight.

### SARS-CoV-2 Neutralization Assay

Production of VSV pseudotyped with SARS-CoV-2 S: 293T cells plated 24 hours previously at 5 × 10^6^ cells per 10 cm dish were transfected using calcium phosphate with 35 μg of pCG1 SARS-CoV-2 S D614G delta18 expression plasmid encoding a codon optimized SARS-CoV2 S gene with an 18-residue truncation in the cytoplasmic tail (kindly provided by Stefan Pohlmann). Twelve hours post transfection the cells were fed with fresh media containing 5mM sodium butyrate to increase expression of the transfected DNA. Thirty hours after transfection, the SARS-CoV-2 spike expressing cells were infected for 2-4 hours with VSV-G pseudotyped VSVΔG-RFP at an MOI of ^∼^1-3. After infection, the cells were washed twice with media to remove unbound virus. Media containing the VSVΔG-RFP SARS-CoV-2 pseudotypes was harvested 28-30 hours after infection and clarified by centrifugation twice at 6000g then aliquoted and stored at −80 °C until used for antibody neutralization analysis.

Antibody neutralization assay using VSVΔG-RFP SARS-CoV-2: All sera were heat-inactivated for 30 minutes at 55 °C prior to use in neutralization assay. Vero E6 cells stably expressing TMPRSS2 were seeded in 100 μl at 2.5×10^4^ cells/well in a 96 well collagen coated plate. The next day, 2-fold serially diluted serum samples were mixed with VSVΔG-RFP SARS-CoV-2 pseudotype virus (100-300 focus forming units/well) and incubated for 1hr at 37 °C. Also included in this mixture to neutralize any potential VSV-G carryover virus was 1E9F9, a mouse anti-VSV Indiana G, at a concentration of 600 ng/ml (Absolute Antibody, <u>Ab01402-2.0</u>). The serum-virus mixture was then used to replace the media on VeroE6 TMPRSS2 cells. 22 hours post infection, the cells were washed and fixed with 4% paraformaldehyde before visualization on an S6 FluoroSpot Analyzer (CTL, Shaker Heights OH). Individual infected foci were enumerated, and the values compared to control wells without antibody. The focus reduction neutralization titer 50% (FRNT_50_) was measured as the greatest serum dilution at which focus count was reduced by at least 50% relative to control cells that were infected with pseudotype virus in the absence of human serum. FRNT_50_ titers for each sample were measured in at least two technical replicates and were reported for each sample as the geometric mean.

### ELISA

Flat bottom, high binding polystyrene 96-well plates (Corning, 9018) were coated overnight at 4°C with either 2 ug/ml SARS-CoV-2 Spike Protein ECD (Sino Biological, 40589-V08B1) or 1 ug/ml SARS-CoV-2 Spike-RBD (Sino Biological, 40592-V08B-B) in PBS. The following day the coating buffer was aspirated, and the plate was washed with PBST and then dried. Once dried the plate was blocked with PBS + 1% BSA for 2 hr at RT before heat inactivated serum samples diluted in PBS + 1% BSA were added for 1 hour at room temperature. The ELISA plate was then washed 4 times with PBS + 0.05% Tween-20 (PBST), followed by addition of 100 uL of 1:2000 goat anti-mouse IgG-ALP (Abcam, ab97020) diluted in PBST + 1% BSA. After 1 hour at room temperature, plates were washed 4 times with PBST and then developed with PNPP (Thermo, 37621) for 30 minutes at room temperature. Optical density (OD) was measured at 405 nm. Endpoint titers were calculated as the highest reciprocal dilution that emitted an optical density exceeding 3× secondary antibody only control.

### Mouse ELISpots

Mouse IL-2 (MABTECH, 3441-4APW-2), IL-4 (MABTECH, 3311-4APW-2), and IL-10 (MABTECH, 3432-4APW-2) ELISpotPLUS plates as well as Mouse IFNγ ELISpotPRO plates (MABTECH, 3421M-2AST-2) were washed with PBS and blocked overnight at 4°C in RPMI 1640 media (Invitrogen, 11875085) + 10 % FBS (Life Technologies, 16000044) + Antibiotic-Antimycotic (Invitrogen, 15240062) + GlutaMAX (Invitrogen, 35050061) (RPMI10+ media). The next day, spleens were processed into single cell suspensions in PBS and then spun at 1500g for 10 min. Red blood cells were lysed with ACK lysis buffer (Quality Biological Inc., 118-156-721EA) + 1:500 benzonase (Sigma, E1014-25KU) for 10 minutes at RT. Splenocytes were plated in RPMI10+ media at 2E5 cells per well and stimulated for 20 hrs (37°C, 5% CO2) with peptide pools of 15-mers with 11 amino acid overlap covering either the S1 or S2 fragment of either WA1 SARS-CoV-2 spike protein (JPT, PM-WCPV-S-1) or B.1.1.7/alpha variant spike protein (JPT, PM-SARS2-SMUT01-1) at a final concentration of 5 ug/ml. PMA/Ionomycin was used as a positive control (eBioscience, 00-4970-03) for all murine ELISpots and RPMI10+ media alone was used as a negative control for each animal. Plates were then incubated in ddH_2_O for 2 minutes and then washed 5× with PBS before incubating with biotinylated anti-cytokine antibody at the manufacture’s recommended concentration in PBS + 1% BSA for 2 hr at RT. Following another 5× PBS wash plates were incubated with Streptavidin-ALP in PBS + 1% BSA for 1 hr at RT and then 5× washed again with PBS. Nitor-blue Tetrazolium Chloride/5-bromo-4-chloro 3 ‘indolyl phosphate p-toludine salt (NBT/BCIP chromagen) substrate solution was added after the final wash for 10-12 min to allow spots to form. Spots were quantified on a CTL ImmunoSpot S6 Core Analyzer.

### Macaques

All procedures were approved by the Institutional Animal Care and Use Committee of the Children’s Hospital of Philadelphia (CHOP). Animals were negative for viral pathogens including SIV (simian immunodeficiency virus), STLV (simian-T-lymphotrophic virus), SRV (simian retrovirus), and B virus (herpesvirus 1) and housed in an AAALAC-accredited facility at CHOP on a 12-hour timed light/dark cycle. Animals received varied enrichment including food treats, manipulatives, visual and auditory stimuli, and social interactions throughout the study. Four 2-3yo female cynomolgus monkeys (*Macaca fascicularis*) were treated with the clinical candidates intramuscularly at doses of 1E11 or 1E12 gc/animal. Serum and PBMC samples were obtained at baseline and 2-week intervals for analyses of immunogenicity. At necropsy, splenocytes, popliteal and inguinal lymph nodes and exposed and unexposed skeletal muscle were harvested for immunologic and histopathologic studies.

### Macaque Immunohistochemistry

Skeletal muscle harvested at necropsy was embedded in O.C.T. (Tissue-Tek, 4583) within 7×7×5 mm base molds (Richard-Allan Scientific, 58949) by rapid submersion in liquid nitrogen-cooled isopentane. All steps with tissue sectioning and IHC are the same as with mouse tissue through blocking in 10% normal donkey serum (Abcam, ab7475) for 20 minutes at RT. After blocking, slides were incubated in PBS with rat anti-CD8 (Bio-Rad, YTC182.20) plus either rabbit polyclonal anti-SARS-CoV-2 Spike/RBD (Sino, 40592-T62) or rabbit polyclonal anti-CD11b (Abcam, ab52478) for 1 hour at 37°C. Slides were then washed 3×10 minutes in PBS at RT and again blocked with 10% normal donkey serum (Abcam, ab7475) for 20 minutes at room temperature before being incubated with PBS containing WGA-555 (Thermo Fisher, W32464), donkey anti-rabbit-AF488 (Abcam, ab150061), and donkey anti-rat-AF647 (Abcam, ab150155) for 30 minutes at 37°C. Following another 3×10 min washes in PBS, slides were mounted in VECTASHIELD mounting media containing DAPI (Vector Laboratories, H-1500). Images were taken using a Zeiss Observer 7 widefield microscope (Zeiss).

### Macaque ELISpot

Monkey IFNγ ELISpotPRO plates (MABTECH, 3421M-2AST-2) were washed with PBS and blocked overnight at 4°C in RPMI10+ media. 2E5 macaque PBMCs were plated in RPMI10+ media and stimulated for 24 hrs (37°C, 5% CO2) with peptide pools of 15-mers with 11 amino acid overlap covering either the S1 or S2 fragment of WA1 SARS-CoV-2 spike protein (JPT, PM-WCPV-S-1) at a final concentration of 5 ug/ml. PMA/Ionomycin was used as a positive control (eBioscience, 00-4970-03) and RPMI10+ media alone was used as a negative control for each animal. Plates were then incubated in ddH_2_O for 2 minutes and then washed 5× with PBS before incubating with ALP-conjugated anti-IFNγ antibody at the manufacture’s recommended concentration in PBS + 1% BSA for 2 hr at RT. Following 5× washes in PBS NBT/BCIP chromagen substrate solution was added for 15 min to allow spots to form. Spots were quantified on a CTL ImmunoSpot S6 Core Analyzer.

## Data availability

All data generated or analyzed during this study are included in this published article or in the Supplementary Information files.

---

**Extended Data Figure 1:**
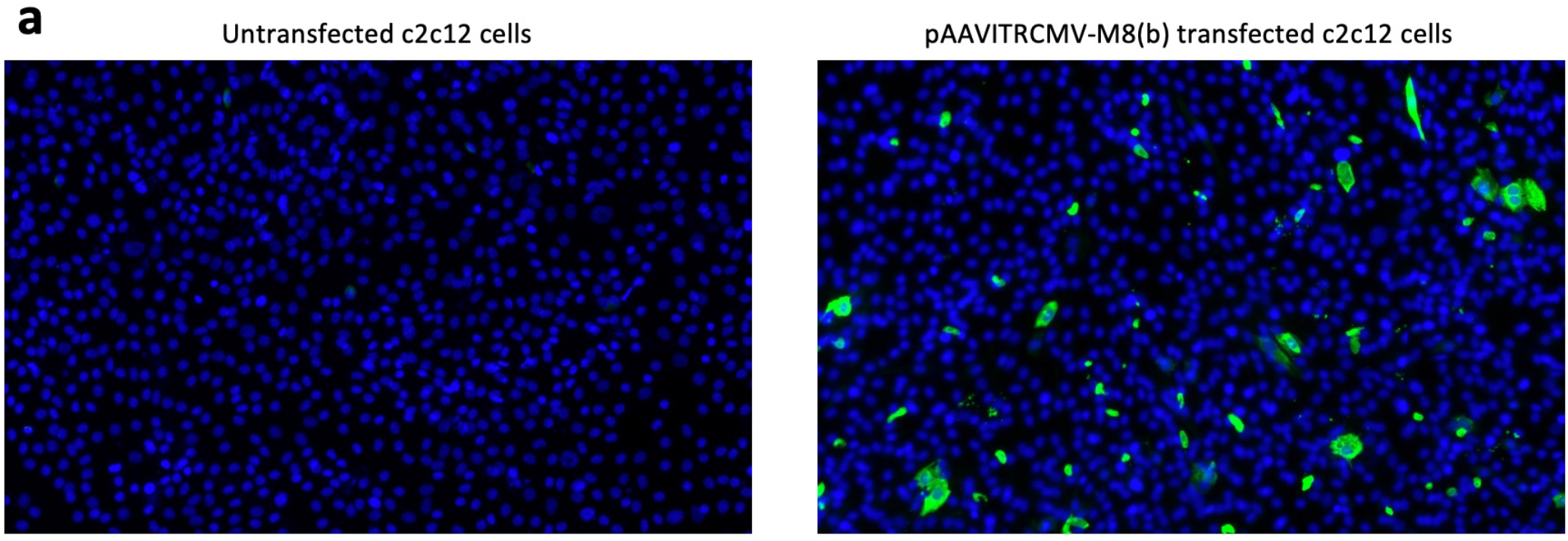
*In vitro* characterization. **a, c**2c12 cells were lipofectamine transfected with plasmid harboring the following elements in sequence AAV_ITR-CMV-UTR5’-M8B-3’UTR-pA-AAV_ITR and then grown on chamber slides. Later cells were fixed and stained for spike protein and nuclei labeled with DAPI.

**Extended Data Figure 2:**
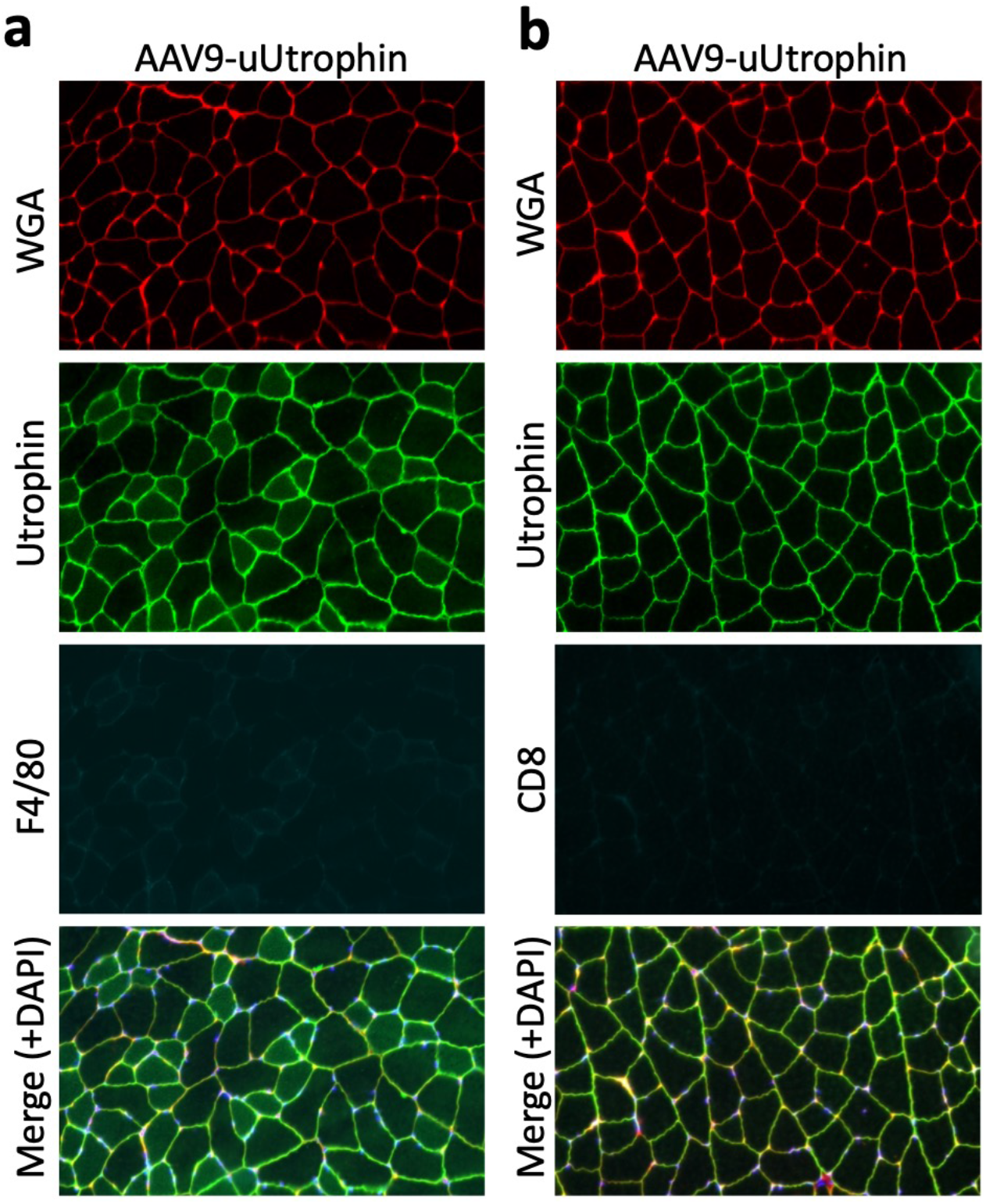
AAV vaccine immunogenicity is directed against foreign M8/M8B spike protein, not AAV capsid. F4/80+ macrophages and CD8+ cytotoxic T-cells fail to infiltrate and surround ⍰Utrophin expressing muscle fibers at 3 weeks post 50 ul IM injection of 6.4E10 vg. AAV9-uUtrophin.

**Extended Data Figure 3:**
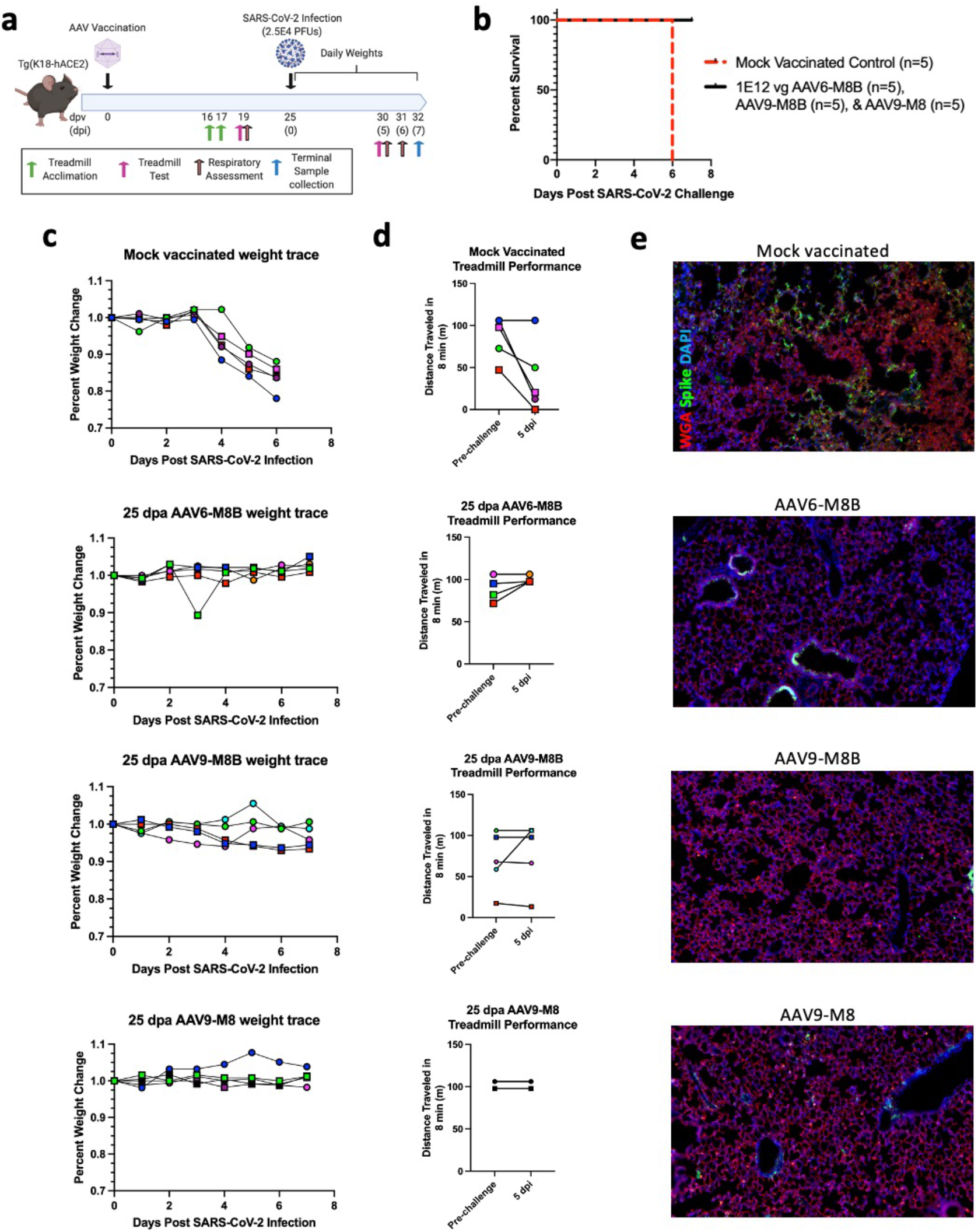
Protection and efficacy in k18-hACE2 mice. **a**, Experimental schematic for data depicted in Fig. 3a-g and Extended Data Figure 3. **b,** Kaplan Meier survival curve. **c,** Individual animal weight traces for each animal following 2.5E4 PFU SARS-CoV-2 IN inoculation. **d,** Individual results of the 8-minute treadmill performance test depicted in Fig. 3b. Because there is a maximum performance on the time restricted protocol, if multiple animals achieve a maximal distance, they appear on top of each other. All animals are represented in the graphic. **e,** Lung IHC post SARS-CoV-2 challenge of mice depicted in the schematic in (**a**) for spike protein, WGA (extracellular matrix stain to assess gross lung morphology), and DAPI. (**C-D**) Circles represent females and squares represent males.

**Extended Data Figure 4:**
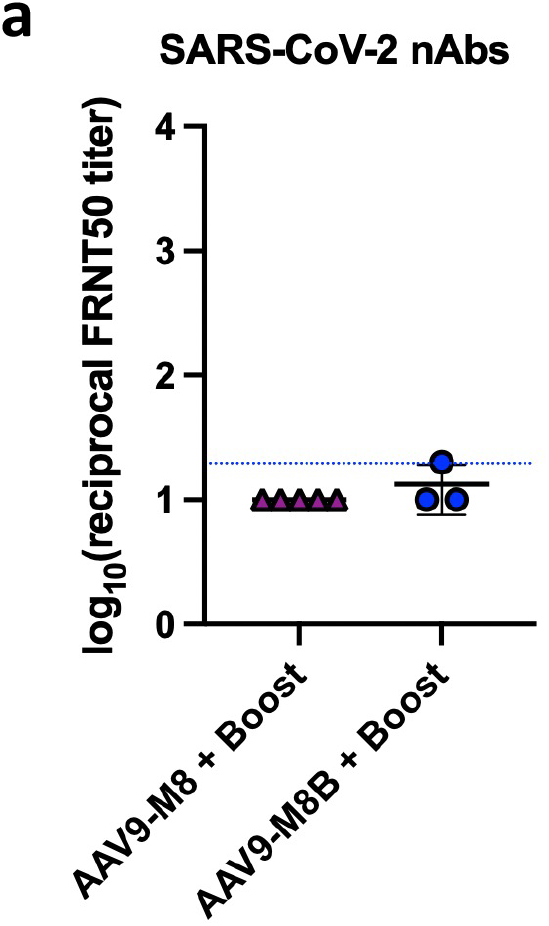
Boosting fails to elicit a neutralizing humoral following booster vaccination. **a,** 2–5-month-old C57BL/10 mice received 6.4E10 vg IM injection of either AAV9-M8 (n=5) or AAV9-M8B (n=3) and then received an identical booster vaccination 5 weeks later. Three weeks after the booster dose animals were evaluated for the presence of SARS-CoV-2 neutralizing antibodies. Data is reported as log_10_(inverse SARS-CoV-2 50% pseudovirus-neutralization titer as measured by focus reduction neutralization test (FRNT_50_)).

**Extended Data Figure 5:**
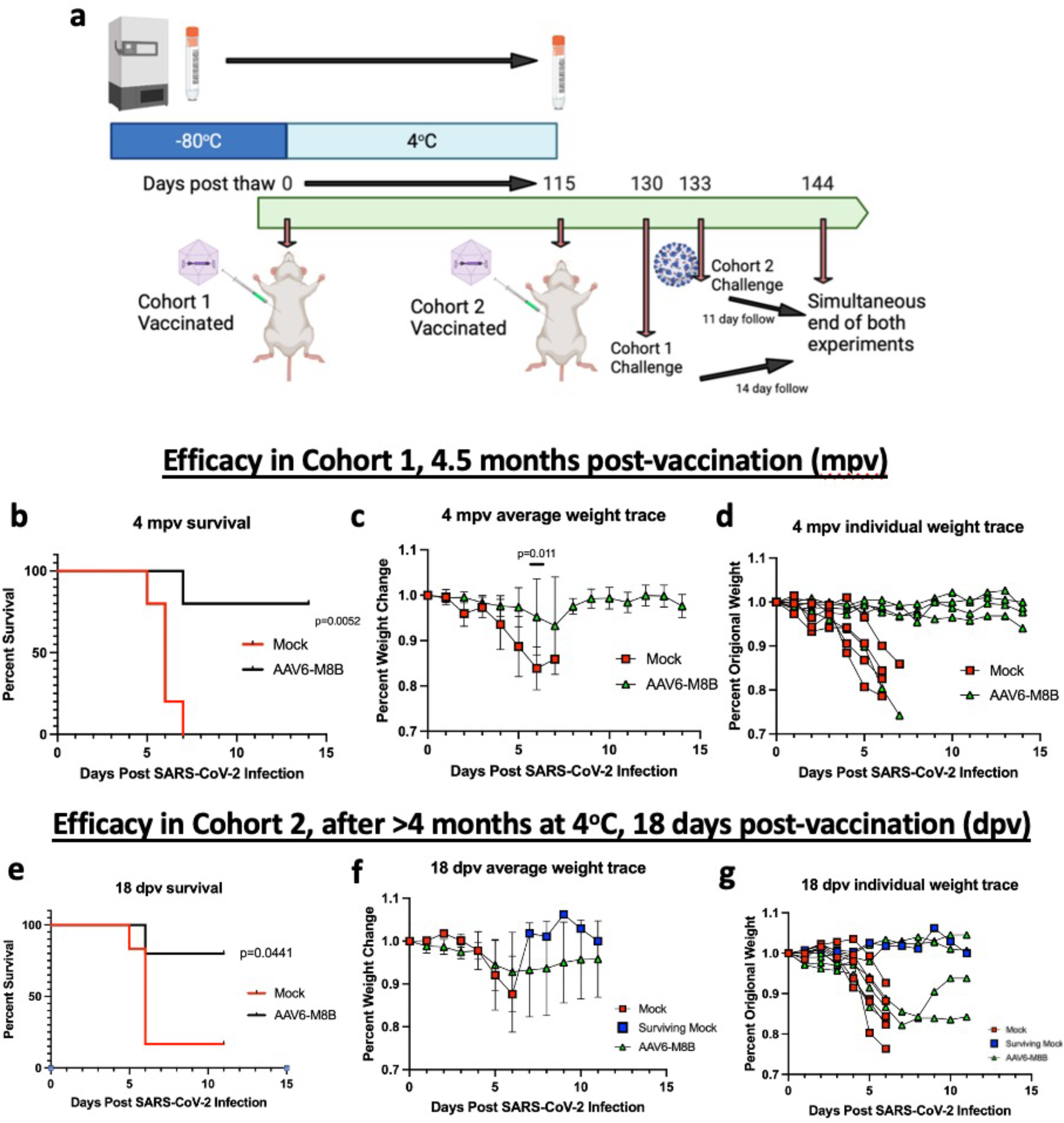
Long term efficacy studies of AAV6-M8B in k18-hACE2 mice. **a,** Schematic of experimental plan. A vial of vector was thawed and immediately used to vaccinate “cohort 1” of k18-hACE2 mice (n=5 AAV6-M8B, n=5 mock). The tube was then kept in at 4°C for >4 months before it was used to vaccinate a second cohort, “cohort 2,” of k18-hACE2 mice (n=5 AAV6-M8B, n=6 mock). Both cohorts were challenged days apart from each other with IN inoculation of 2.5E4 PFU SARS-CoV-2, at 4.5 mpv for cohort 1 (**b-d**) and at 18 dpv for cohort 2 (**e-g**). All animals were tracked until a terminal concluding date. **b, e,** Kaplan Meier survival curve post SARS-CoV-2 infection. Survival was compared by the log-ranked Mantel-Cox test. **c, f,** Change in body weight post SARS-CoV-2 infection. Weight was compared by the nonparametric Mann–Whitney U-test. The average mock vaccinated body weight in (**f**) switches from red to blue when the entire average weight is derived from a single surviving animal. **d, g,** Individual animal traces for animals depicted in (**c, f**).

